# RNA-directed DNA methylation prevents rapid and heritable reversal of transposon silencing under heat stress in Zea *mays*

**DOI:** 10.1101/2021.01.08.425849

**Authors:** Wei Guo, Dafang Wang, Damon Lisch

**Affiliations:** Department of Botany and Plant Pathology, Purdue University, West Lafayette, Indiana, USA; Division of Math and Sciences, Delta State University, Cleveland, MS, USA

## Abstract

In large complex plant genomes, RNA-directed DNA methylation (RdDM) ensures that epigenetic silencing is maintained at the boundary between genes and flanking transposable elements. In maize, RdDM is dependent on *Modifer of Paramutation 1* (*Mop1*), a putative RNA dependent RNA polymerase. Here we show that although RdDM is essential for the maintenance of DNA methylation of a silenced *MuDR* transposon in maize, a loss of that methylation does not result in a restoration of activity. Instead, heritable maintenance of silencing is maintained by histone modifications. At one terminal inverted repeat (TIR) of this element, heritable silencing is mediated via H3K9 and H3K27 dimethylation, even in the absence of DNA methylation. At the second TIR, heritable silencing is mediated by H3K27 trimethylation, a mark normally associated with somatically inherited gene silencing. We find that a brief exposure of high temperature in a *mop1* mutant rapidly reverses both of these modifications in conjunction with a loss of transcriptional silencing. These reversals are heritable, even in *mop1* wild type progeny in which methylation is restored at both TIRs. These observations suggest that DNA methylation is neither necessary to maintain silencing, nor is it sufficient to initiate silencing once has been reversed. However, given that heritable reactivation only occurs in a *mop1* mutant background, these observations suggest that DNA methylation is required to buffer the effects of environmental stress on transposable elements.

**Author Summary:** Most plant genomes are mostly transposable elements (TEs), most of which are held in check by modifications of both DNA and histones. The bulk of silenced TEs are associated with methylated DNA and histone H3 lysine 9 demethylation (H3K9me2). In contrast, epigenetically silenced genes are often associated with histone lysine 27 trimethylation (H3K27me3). Although stress can affect each of these modifications, plants are generally competent to rapidly reset them following that stress. Here we demonstrate that although DNA methylation is not required to maintain silencing of the *MuDR* element, it is essential for preventing heat-induced, stable and heritable changes in both H3K9me2 and H3K27me3 at this element, and for concomitant changes in transcriptional activity. These finding suggest that RdDM acts to buffer the effects of heat on silenced transposable elements, and that a loss of DNA methylation under conditions of stress can have profound and long lasting effects on epigenetic silencing in maize.

## Introduction

Transposable elements (TEs) are a ubiquitous feature of all genomes. They survive in large measure because they can out-replicated the rest of the genome [1]. As a consequence of that replication, TE can threaten the integrity of the host genome. In response to this threat, all forms of life have evolved mechanisms by which TEs can be silenced when they are recognized as such and, importantly, maintained in a silenced state over long periods of time, even when the initial trigger for silencing is no longer present [2–4]. Because plant genomes are largely composed of TEs, the majority of plant DNA is maintained in an epigenetically silent state [5]. Because they are the primary target of epigenetic silencing in plants, TEs are an excellent model for understanding the means by which particular DNA sequences are targeted for silencing, and for understanding the means by which silencing can be maintained from one generation to the next [6]. Finally, because TEs have proved to be exquisitely sensitive to a variety of stresses [7–9], they can also teach us a great deal about the relationship between stress and epigenetically encoded memory of stress.

In plants, heritable epigenetic silencing of TEs is almost invariably associated with DNA methylation [10–12]. The vast bulk of TEs in plant genomes are methylated and, with some notable exceptions [13], epigenetically silenced [14, 15]. DNA methylation has a number of features that makes it an appealing mechanism by which silencing can be heritably propagated, either following cell divisions during somatic development, or transgenerationally, from one generation to the next. Because methylation in both the CG and CHG sequence contexts (where H = A, T or G) are symmetrical, information concerning prior DNA methylation can be easily propagated by methylating newly synthesized DNA strands using the parent strand as a template. For CG methylation, this is achieved by reading the methylated cytosine using VARIANT IN METHYLATION 1-3 (VIM1-3) [16, 17] and writing new DNA methylation using the methyl transferase MET1 [18–20]. For CHG, methylation is read indirectly by recognition of H3K9 dimethylation (H3K9me2) by CMT3, which catalyzes methylation of newly synthesized DNA, which in turn triggers methylation of H3K9 [21–23].

Maintenance methylation of most CHH involves RNA-directed DNA methylation (RdDM). The primary signal for *de novo* methylation of newly synthesized DNA from previously methylated DNA sequences is thought to be transcription by DNA POLYMERASE IV (POLIV) of short transcripts from previously methylated templates [24–26]. This results in the production of small RNAs that are tethered to the target DNA by DNA POLYMERASE IV (POLV), which is targeted by SU(VAR)3-9 homologs SUVH2 and SUVH9, which bind to methylated DNA [27]. This in turn triggers *de novo* methylation of newly synthesized DNA strands using the methyl transferases DRMT1/2 [28, 29]. In addition to the RdDM pathway, CHH methylation can also be maintained due to the activity of CHROMOMETHYLASE (CMT2), which, similar to CMT3, works in conjunction with H3K9me2 to methylate non-CG cytosines, particularly in deeply heterochromatic regions of the genome [30]. Finally, because both histones and DNA must be accessible in order to be modified, chromatin remodelers such as DDM1 are also often required for successful maintenance of TE silencing [23, 31]. In plants, effective silencing of TEs requires coordination between DNA methylation and histone modifications [32]. Together, these pathways can in large part explain heritable propagation of both DNA methylation and histone modification of TEs.

In large genomes such as that of maize, much of RdDM activity is focused not on deeply silenced heterochromatin, which is often concentrated in pericentromeric regions, but on regions immediately adjacent to genes, referred to as “CHH islands” because genes in maize are often immediately adjacent to silenced TEs [15, 33]. In maize, mutations in components of the RdDM pathway affect both paramutation and transposon silencing. Mutations in *Modifier of Paramutation 1* (*Mop1*), a homolog of *RNA DEPENDENT RNA POLYMERASE2* (*RDR2*), result in the loss of nearly all 24 nucleotide small RNAs, as well as the CHH methylation that is associated with them [34–36]. Despite this, *mop1* has only minimal effects gene expression in any tissue except the meristem [33, 37], and the plants are largely phenotypically normal. This, along with similar observations in Arabidopsis, has led to the suggestion that the primary role of RdDM is to reinforce boundaries between genes and adjacent TEs, rather than to regulate gene expression [33].

Unlike animals, plants do not experience a global wave of DNA demethylation either in the germinal cells of the gametophyte or the in early embryo [38]. Thus, DNA methylation and associated histone modifications are an attractive mechanism for transgenerationally propagated silencing. Indeed, there is strong evidence that mutants that trigger a global loss of methylation can cause heritable reactivation of previously silenced TEs, although it is worth noting that even in mutants in which the vast majority of DNA methylation has been lost, only a subset of TEs are transcriptionally reactivated [39, 40], and DNA methylation of many TEs can be rapidly reestablished at many loci via RdDM in wild type progenies of mutant plants, suggesting that memory propagated via DNA methylation can be restored due to the presence of small RNAs that can in trigger *de novo* methylation of previously methylated sequences [41, 42].

In contrast to TEs, most genes that are silenced during somatic development in plants are associated with H3K27 trimethylation (H3K27me3), which requires the activity of the polycomb complexes PRC2 and PRC1, which together catalyze H3K27 methylation and facilitate its heritable propagation [43–45]. In plants, H3K27me3 enrichment is generally associated with genes rather than TEs [46, 47]. The most well explored example of this involves epigenetic setting of FLC, a negative regulator of flowering in Arabidopsis [48, 49]. In a process known as vernalization, prolonged exposure to cold results in somatically heritable silencing of this gene, which in turn results in flowering under favorable conditions in the spring. Somatically heritable silencing of FLC is initially triggered by non-coding RNAs, which are involved in recruitment of components of PRC2, which catalyze H3K27 trimethylation, which in turn mediates a somatically heritable silent state [48]. Importantly, H3K27 trimethylation at genes like FLC is erased each generation, both in pollen and in the early embryo [50–52]. The fact that H3K27me3 must be actively reset suggests that in the absence of this resetting, H3K27me3 in plants is competent to mediate transgenerational silencing but is normally prevented from doing so.

Dramatic differences in TE content between even closely related plant species suggest that despite the relative stability of TE silencing under laboratory conditions, TEs frequently escape silencing and proliferate in natural settings [53]. Stress, both biotic and abiotic can often trigger TE transcription and, at least in some cases, transposition [7, 54–57]. Further, there is evidence that the association of TEs and genes can result in *de novo* stress induction of adjacent genes [54, 58, 59].

Because of its dramatic and global effects on both gene expression and protein stability, heat stress has attracted considerable attention, particularly with respect to heritable transmission of TE activity. For both genes and TEs, although heat stress can trigger somatically heritable changes in gene expression, there appear to be a variety of mechanisms to prevent or gradually ameliorate transgenerational transmission of those changes [60, 61]. Thus, for instance, although the Onsen retrotransposon is sensitive to heat, it is only in mutants in the RdDM pathway that transposed elements are transmitted to the next generation [9, 62]. Given that both TEs and various components of regulatory pathways that have evolved to regulated them are up-regulated in germinal lineages, this is not surprising [63, 64]. Similar experiments using silenced transgenes have demonstrated that double mutants of *mom1* and *ddm1* cause these transgenes as well as several TEs to be highly responsive to heat stress, and the observed reversal of silencing can be passed on to a subsequent generation, but only in mutant progeny [65]. It is also worth noting that in many cases of TE reactivation, silencing is rapidly re-established in wild type progeny [66, 67]. The degree to which this is the case likely depends on a variety of factors, from the copy number of a given element, its position within the genome, its mode of transposition and the presence or absence of trans-acting small RNAs targeting that TE [68].

Our model for epigenetic silencing is the *Mutator* system of transposons in maize. The *Mutator* system is a family of related elements that share similar, 200 bp terminal inverted repeats but that contain distinct internal sequences. Nonautonomous *Mu* elements can only transpose in the presence of the autonomous element, *MuDR. MuDR* is a member of the MULE superfamily of Class II cut and paste transposons [69, 70]. In addition to being required for transposition, the 200 bp TIRs within *MuDR* elements serve as promoters for the two genes encoded by *MuDR, mudrA*, which encodes a transposase, and *mudrB*, which encodes a novel protein that is required for *Mu* element integration. Both genes are expressed at high levels in rapidly dividing cells, and expression of both of them is required for full activity of the *Mutator* system [71, 72]. MURA, the protein produced by *mudrA*, is sufficient for somatic excision of *Mu* elements, which results in characteristically small revertant sectors in somatic tissue. *MuDR* elements can be heritably silenced when they are in the presence of *Mu killer* (*Muk*), a rearranged variant of *MuDR* whose transcript forms a hairpin that is processed into 21-22 nt small RNAs that directly trigger transcriptional gene silencing (TGS) of *mudrA* and indirectly trigger silencing of *mudrB* when it is in trans to *mudrA [4, 73]*. Because *Muk* can be used to heritably silence *MuDR* through a simple cross, and because silencing of *MuDR* can be stably maintained after *Muk* is segregated away, the *MuDR/Muk* system is an excellent model for understanding both initiation and maintenance of silencing. Prior to exposure to *Muk, MuDR* is fully active and is not prone to spontaneous silencing [74]. After exposure, *MuDR* silencing is exceptionally stable over multiple generations [73].

When *mudrA* is silenced, DNA methylation in all three sequence contexts accumulates within the 5’ end of the TIR immediately adjacent to *mudrA* (TIRA) [75]. Methylation at the 5’ and 3’ portions of this TIR have distinctive causes and consequences. The 5’ end of the TIR is readily methylated in the absence of the transposase, but this methylation does not induce transcriptional silencing of *mudrA* [76]. Methylation in this end of TIRA is readily eliminated in the presence of functional transposase. However, the loss of methylation in a silenced element in this part of the TIRA does not result in heritable reactivation of a silenced element. In contrast, CG and CHG methylation the 3’ portion of TIRA, which corresponds to the *mudrA* transcript as well as to *Muk*-derived 22 nt small RNAs that trigger silencing, is not eliminated in the presence of active transposase and is specifically associated with heritable transcriptional silencing of *mudrA*.

The second gene encoded by *MuDR* elements, *mudrB*, is also silenced by *Muk*, but the trajectory of silencing of this gene is entirely distinct, despite the fact that the *Muk* hairpin has near sequence identity to the TIR adjacent to *mudrB* (TIRB) [4, 73]. By the immature ear stage of growth in F1 plants that carry both *MuDR* and *Muk*, *mudrA* is transcriptionally silenced and densely methylated. In contrast, *mudrB* in intact elements remains transcriptionally active in this tissue, but its transcript is not polyadenylated. It is only in the next generation that steady state levels of transcript become undetectable. Further, experiments using deletion derivatives of *MuDR* that carry only *mudrB* are not silenced by *Muk* when they are on their own, or when they are in trans to an intact *MuDR* element that is being silenced by *Muk*. This suggests that heritable silencing of *mudrB* is triggered by the small RNAs that target *mudrA*, but the means by which this occurs is indirect and involves spreading of silencing information from *mudrA* to *mudrB*.

Silencing of *mudrA* can be destabilized by the *mop1* mutant, a homolog of RNA-DEPENDENT RNA POLYMERASE2 (RDR2) that is required for the production of the vast bulk of 24 nt small RNAs in maize, including those targeting *Mu* TIRs [34–36, 77]. However silencing of *MuDR* by *Muk* is unimpeded in a *mop1* mutant background, likely because *Muk*-derived small RNAs are not dependent on *mop1 [78]*. Further, although reversal of silencing of *MuDR* in a *mop1* mutant background does occur, it only occurs gradually, over multiple generations, and only affects *mudrA*. In contrast, *mudrB* is not reactivated in this mutant background and, because *mudrB* is required for insertional activity, although these reactivated elements can excise during somatic development, they cannot insert into new positions.

## Results

### DNA methylation is not required to maintain silencing of *MuDR* elements in *mop1* mutants

Given that *MuDR* elements are only activated after multiple generations in a *mop1* mutant background, we wanted to understand how silencing of *MuDR* is maintained in *mop1* mutants prior to reactivation. To do this, we examined expression and DNA methylation at TIRA by performing bisulfite sequencing of TIRA of individuals in families that were segregating for a single silenced *MuDR* element, designated *MuDR\**, and that were homozygous or heterozygous for *mop1* (Fig 1A and Fig S1).

**Figure 1.**
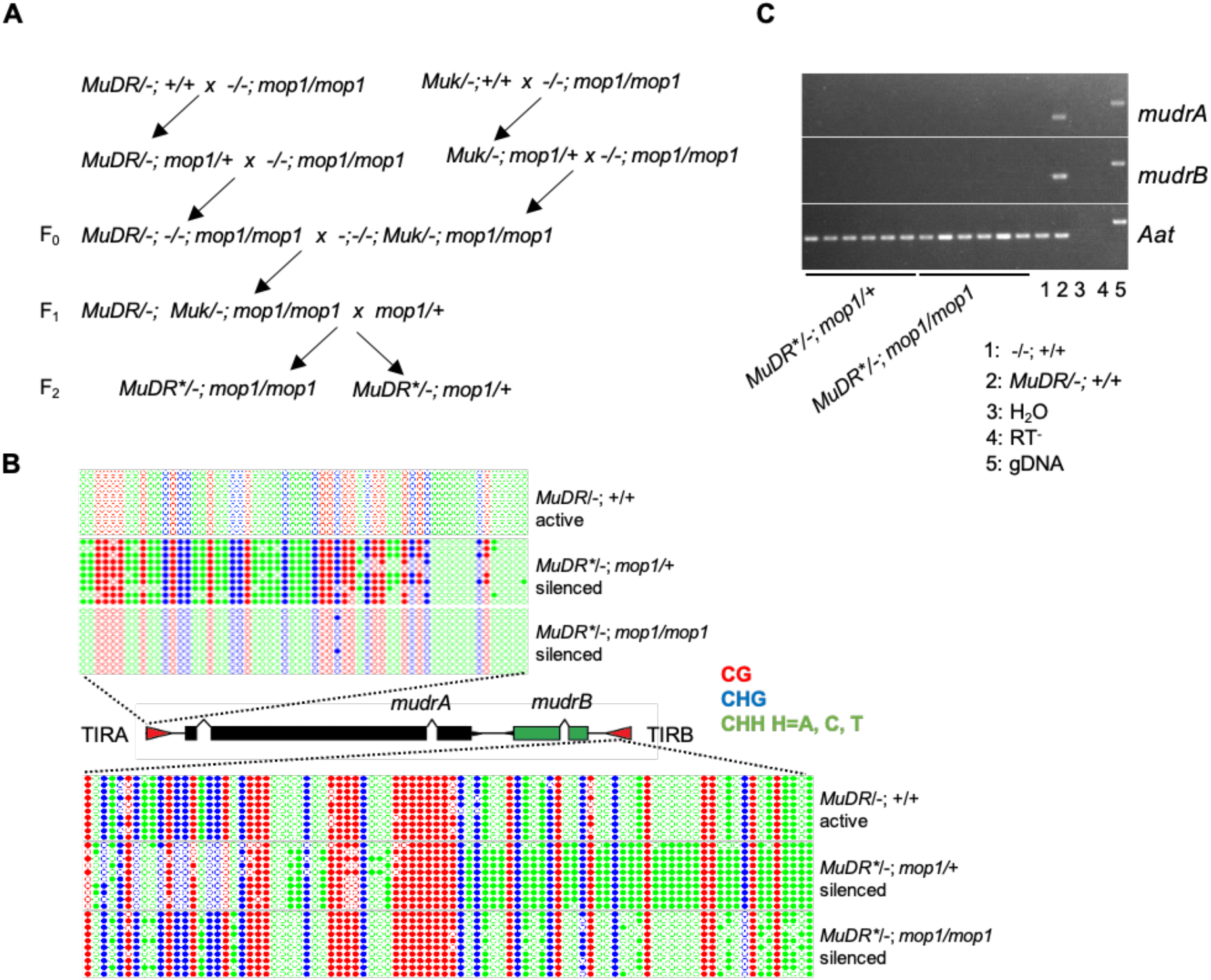
DNA methylation patterns at TIRA and TIRB of stably silenced F2 plants. (A) Crosses used to generate the materials analyzed. (B) DNA methylation patterns at TIRA and TIRB. Ten individual clones were sequenced from amplification of bisulfite-treated samples of the indicated genotypes. The cytosines in different sequence contexts are represented by different colors (red, CG; blue, CHG; green, CHH, where H=A, C, or T). For each genotype, DNA from six biological replicates were pooled. (C) RT-PCR detecting *mudrA* and *mudrB* transcripts in F_2_ plants from a family segregating for a single silenced *MuDR* element (*MuDR*), mop1/+ and mop1/mop1*. H_2_O: water control. RT^-^: no reverse transcriptase added. gDNA: genomic DNA.

In control plants carrying an active *MuDR* element, all cytosines in TIRA were unmethylated, which was consistent with our previous results and which indicated that bisulfite conversion was efficient (Fig 1B). Also consistent with previous results, F_2_ *MuDR*/−; mop1/+* plants, whose F_1_ parent carried both *MuDR* and *Muk*, exhibited dense methylation at TIRA. In contrast, DNA methylation in the CG, CHH and CHG contexts at TIRA was absent in *mop1* mutant siblings. Interestingly, *mop1* had effects on TIRB that are more consistent with the known effects of this mutant specifically on CHH methylation. While F_2_ *MuDR*/−; mop1/+* plants exhibited dense methylation at TIRB in all sequence contexts, *mop1* homozygous siblings exhibited a loss of methylation only in the CHH context. Despite the effects of *mop1* on *MuDR* methylation at both TIRA and TIRB, RT-PCR results demonstrated that these *mop1* mutant plants did not exhibit reactivation of *mudrA* or *mudrB* (Fig 1C).

### *MOP1* enhances enrichment of H3K9 and H3K27 dimethylation at TIRA

Transposon silencing is often associated with H3K9 and H3K27 dimethylation, two hallmarks of transcriptional silencing in plants [21, 47]. DNA methylation, particularly in the CHG context, is linked with H3K9 dimethylation through a self-reinforcing loop, and these two epigenetic marks often colocalize at TEs and associated nearby genes [79]. We had previously demonstrated that these two repressive histone modifications corresponded well with DNA methylation of silenced *MuDR* elements at TIRA [75]. However, our observation that silencing of *mudrA* can be maintained in the absence of DNA methylation in *mop1* mutants suggests that additional repressive histone modifications may be responsible for maintaining the silenced state of *mudrA*. To test this hypothesis, we examined the enrichment of H3K9me2 at TIRA in individuals in a family that segregated for silenced *MuDR* and for *mop1* homozygotes and heterozygotes (Fig 1A) by performing a chromatin immunoprecipitation quantitative PCR (ChIP-qPCR) assay. As controls, we also examined these two histone modifications in leaf tissue from plants carrying active and deeply silenced *MuDR* elements in a wild type background. Compared with active *MuDR/−; +/+* plants, H3K9me2 and levels were significantly enriched at TIRA in the *MuDR\**/−; +/+ plants (Fig 2A). The same was true of H3K27me2 (Fig S2). Surprisingly, a significant increase in H3K9me2 and H3K27me2 at TIRA was observed in *mop1* mutants compared with their sibling *mop1* heterozygous siblings and with the silenced *MuDR*/−;* +/+ control plants, suggesting that the loss of DNA methylation that resulted from the loss of MOP1 in these mutants actually resulted in an increase in both of these repressive chromatin marks.

**Figure 2.**
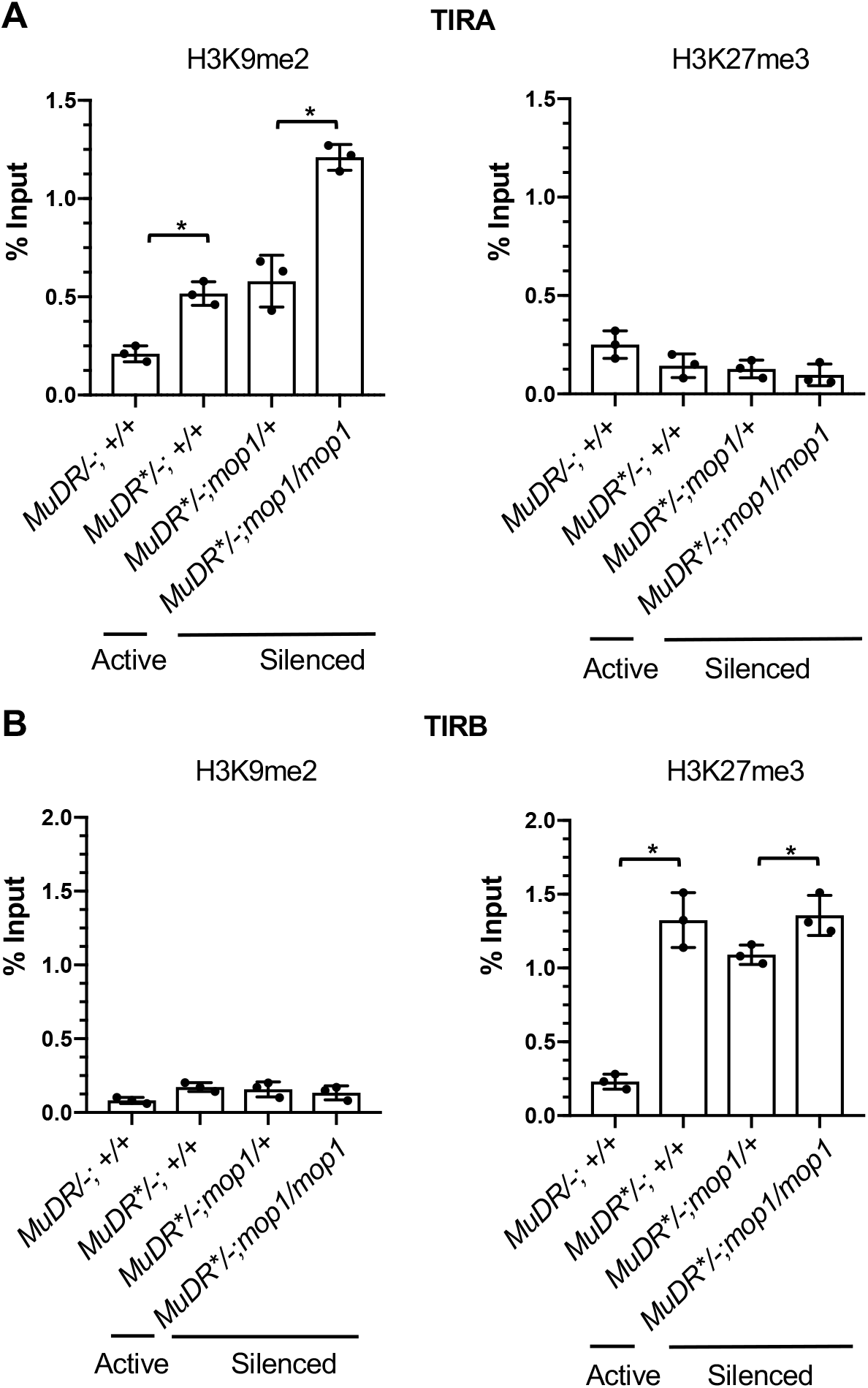
ChIP-qPCR analysis of enrichment of histone marks H3K9me2 and H3K27me3 at TIRA and TIRB in *mop1* mutants. ChIP-qPCR analysis of enrichment of histone marks, H3K9me2 and H3K27me3 at TIRA and TIRB. (A) Relative enrichment of H3K9me2 and H3K27me3 in leaf 3 of plants of the indicated genotypes.*MuDR*: active element.*MuDR\**: inactive element. (B) Relative enrichment of H3K9me2 and H3K27me3 in leaf 3 of plants of the indicated genotypes. qPCR signal was normalized to *Copia* and then to the value of input sample. All data are the average of two technical replicates from three independent lines. An unpaired t-test was performed. Error bars indicate mean ± standard deviation (SD) of the three biological replicates. *P<0.05; **P < 0.01

### Silencing of TIRB is associated with an increase in H3K27me3

Like *mudrA*, *mudrB* is silenced by *Muk*, but maintenance of *mudrB* silencing has distinct requirements. Unlike *mudrA*, which is eventually reactivated in a *mop1* mutant background under normal conditions, *mudrB* remains silenced, suggesting that maintenance of silencing of this gene is independent of *MOP1* [35]. ChIP-qPCR revealed that silencing of *mudrB* is not associated with H3K9me2 methylation. Instead, heritably silenced TIRB is enriched for H3K27me3, a modification normally associated with somatically silenced genes rather than transposable elements (Fig 2B). The *mop1* mutant appears to enhance H3K27me3 at TIRB relative to the *mop1* heterozygous siblings, although the enrichment is no greater that observed in the *MuDR*/−;* +/+ controls.

### Application of heat stress specifically in the early stage of growth can promote the reactivation of silenced *MuDR* elements in *mop1* mutants

There is ample evidence that a variety of stresses can reactivate epigenetically silenced TEs. One particularly effective treatment is heat stress. Given that a loss of methylation by itself is not sufficient to reactivate silenced *MuDR* elements, we subjected *mop1* mutant and *mop1* heterozygous sibling seedlings carrying silenced *MuDR* elements (*MuDR\**) to heat stress. Fourteen-day-old *MuDR*/−; mop1/mop1* and *MuDR*/−; mop1/+* sibling seedlings were heated at 42 °C for four hours and leaf samples were collected immediately after that treatment (Fig 3A). RT-PCR for the heat response factor Hsp90 (Zm00001d024903) confirmed that the seedlings were responding to the heat treatment (Fig S3). We then examined *MuDR* transcription by performing RT-PCR on RNA from leaf three immediately after the plants had been removed from heat and from control plants that had not been subjected to heat stress. In the *mop1* mutants, both *mudrA* and *mudrB* became transcriptionally reactivated upon heat treatment (Fig 3B). *MuDR* elements in plants that were *mop1* mutant that were not heat stressed and were those that were wild type and that were heat stressed were not reactivated, demonstrating that both a mutant background and heat stress are required for efficient reactivation. To determine if the application of heat stress at a later stage of plant development can also promote reactivation, we heat-stressed 28-day-old plants and examined *MuDR* transcription in leaf seven at a similar stage of development (~10 cm) as had been examined in heat stressed leaf three in the previous experiment. In these plants, we saw no evidence of reactivation, indicating that *MuDR* responsiveness to heat shifts over developmental time (Fig 3C). Taken together, these data suggest that the application of heat stress specifically at an early stage of plant development can promote the reactivation of a silenced TE in a mutant that is deficient in the RdDM pathway.

**Figure 3.**
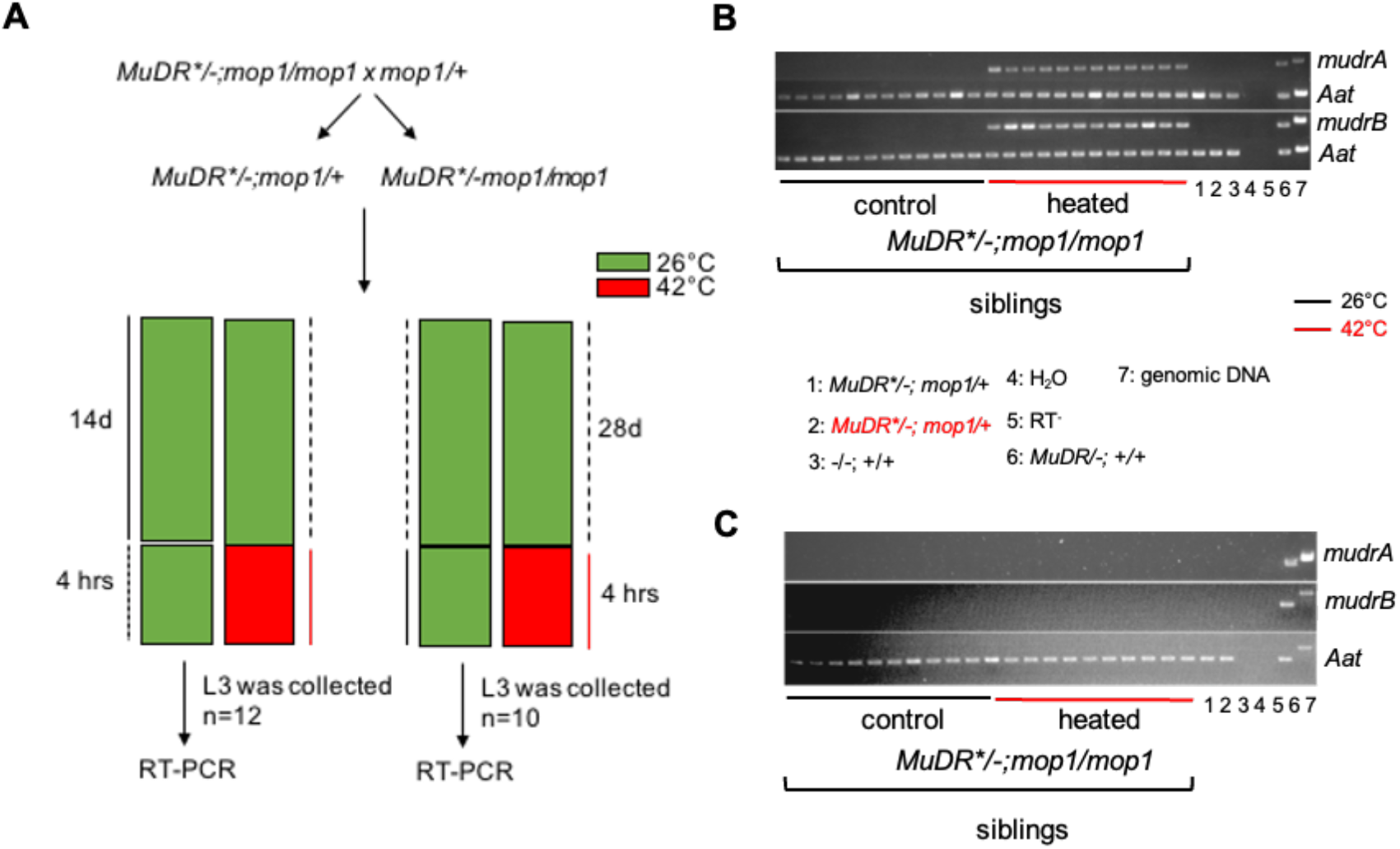
Expression of *mudrA* and *mudrB* in plants under heat stress. (A) Schematic diagram of the heat-reactivation experiment. (B) RT-PCR of *mudrA* and *mudrB* in plants of the indicated genotypes. (C) RT-PCR of *mudrA* and *mudrB* of leaf 7 of heat-treated F2 plants. *Aat* is a housekeeping gene that was used as a positive expression control. Additional controls for each experiment included pools of ten *MuDR/−; mop1/+* heated and ten unheated plants, as well as plants that lacked *MuDR* and were wild type for *mop1 (−/−; +/+)*, samples with water or with no reverse transcriptase as negative controls, active *MuDR* as well as genomic DNA (gDNA) as positive controls for the *MuDR*-specific PCR primers.

TIRA in a *mop1* mutant background already lacks any DNA methylation prior to heat treatment and thus heat would not be expected to reduce TIRA methylation. However in *mop1* mutants TIRB retained CG and CHG methylation and also remained inactive (Fig 1B). To determine if reactivation after heat treatment is associated with a loss of this methylation, we examined DNA methylation at TIRB in *mop1* mutants in the presence or absence of heat treatment. This assay was performed on the same tissues that we collected for *MuDR* expression reactivation analysis. We found that the DNA methylation pattern was the same for both the heat treated and the control *mop1* mutant plants, indicating that heat stress does not alter TIRB methylation and that a further loss of DNA methylation is not the cause of *mudrB* reactivation in this tissue (Fig S4).

### Heat stress reverses TE silencing by affecting histone modifications at TIRA and TIRB

Under normal conditions, we found that H3K9me2 at TIRA is associated with silencing, and H3K9me2 is actually enriched when TIRA methylation is lost in *mop1* mutants (Fig 2A). In contrast, we find that H3K27me3, rather than H3K9me2, is enriched at TIRB and is maintained at similar or slightly elevated levels in *mop1* mutant relative to *mop1* heterozygous siblings (Fig 2B). Given these observations, we hypothesized that heat stress may reverse H3K9me2 enrichment at TIRA and H3K27me3 enrichment at TIRB. To test this hypothesis, we determined the level of H3K9me2 and H3K27me3 at TIRA and TIRB under normal and stressed conditions using ChIP-qPCR.

Upon heat stress, the level of H3K9me2 at TIRA was significantly decreased in *mop1* mutants compared to that of non-treated *mop1/mop1* mutant siblings (Fig 4A). Interestingly, however, H3K9me2 enrichment only decreased to the level observed at TIRA in silenced *MuDR*/−;* +/+ plants, and it remained significantly higher than that of TIRA in the naturally active *MuDR/−; +/+* plants. In contrast, we observed no changes in H3K27me3 at TIRA.

**Figure 4.**
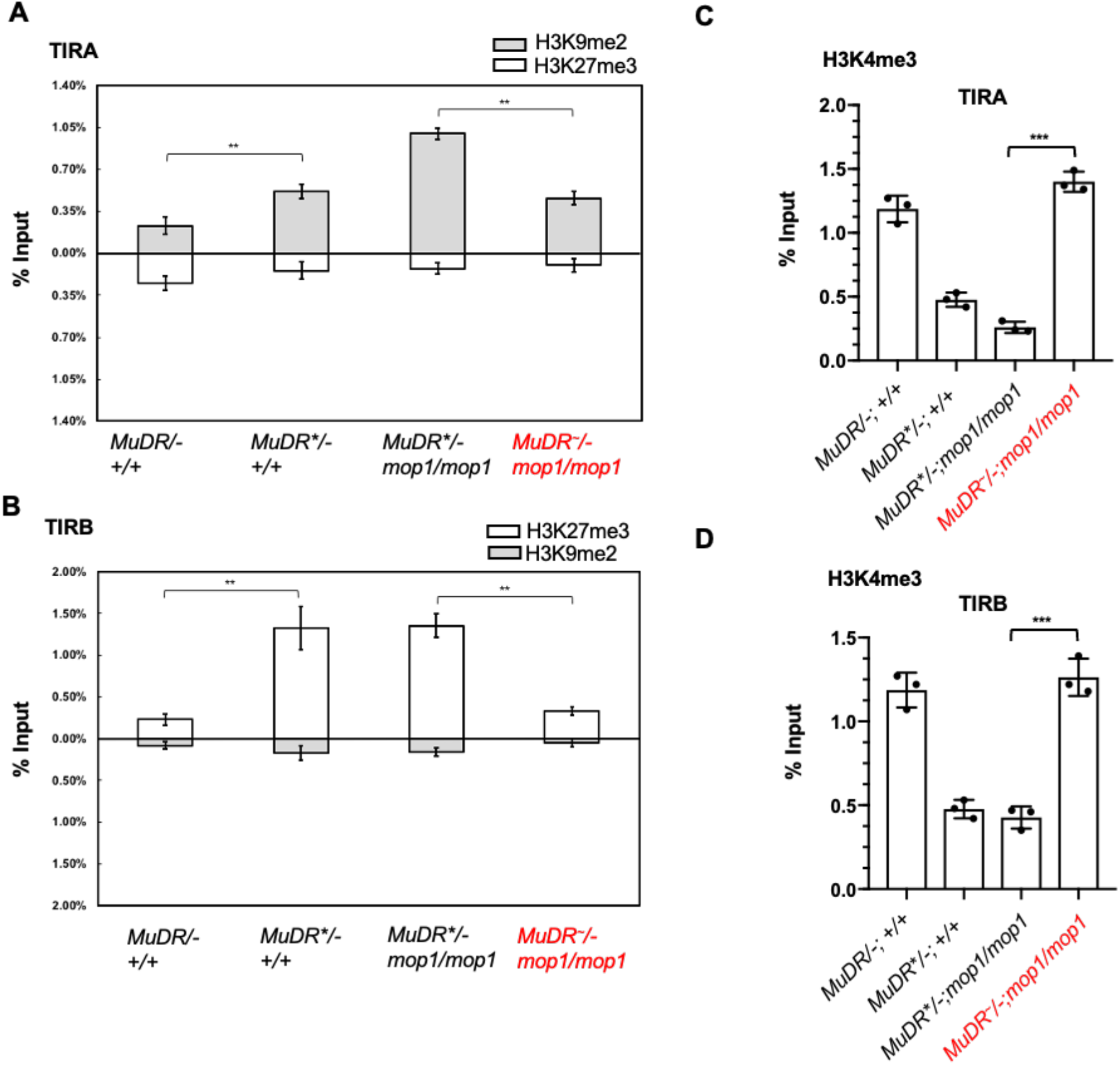
ChIP-qPCR analysis of histone marks TIRA and TIRB under heat stress. Relative enrichment of H3K9me2 and H3K27me3 at TIRA (A) and TIRB (B) in leaf 3 of plants of the indicated genotypes. (Relative enrichment of H3K4me3 at TIRA (C) and TIRB (D) in leaf 3 of plants of the indicated genotypes. qPCR signals were normalized to *Copia* and then to the value of input samples. All data are the average of two technical replicates from three independent lines. An unpaired t-test was performed. Error bars indicate mean ± standard deviation (SD) of the three biological replicates. **P < 0.01; ***P < 0.001

At TIRB, we observed no changes in H3K9me2 enrichment in any of our samples. Instead, we found that heat treatment reversed previously established H3K27me3 at TIRB, supporting the hypothesis that this modification, rather than H3K9me2, mediates heritable silencing of *mudrB* (Fig 4B). Consistent with evidence for transcriptional activation of both *mudrA* and *mudrB*, we observed enrichment of the active mark H3K4me3 in reactivated TIRA and TIRB (Fig 4C,D). Taken together, these data demonstrate that heat stress can simultaneously reduce two often mutually exclusive repressive histone modifications, H3K9me2 and H3K27me3 at the two ends of a single TE.

### The reactivation state is somatically transmitted to the new emerging tissues

We next sought to determine whether or not the reactivated state can be propagated to cells in somatic tissues after the heat had been removed. We performed quantitative RT-PCR to detect *mudrA* and *mudrB* transcripts in mature leaf ten of plants 35 days after the heat stress and in immature tassels ten days after that. At V2, when the heat stress was applied and leaf three was assayed, cells within leaf 10 primordia are present and may have experienced the heat stress. In contrast, because the tassel primordia are not formed until V5, the cells of the tassel could not have experienced the heat stress directly [80, 81]. We found that both genes stayed active in both tissues, indicating heat-induced reactivation is stably transmitted to new emerging cells and tissues (Fig 5).

**Fig 5.**
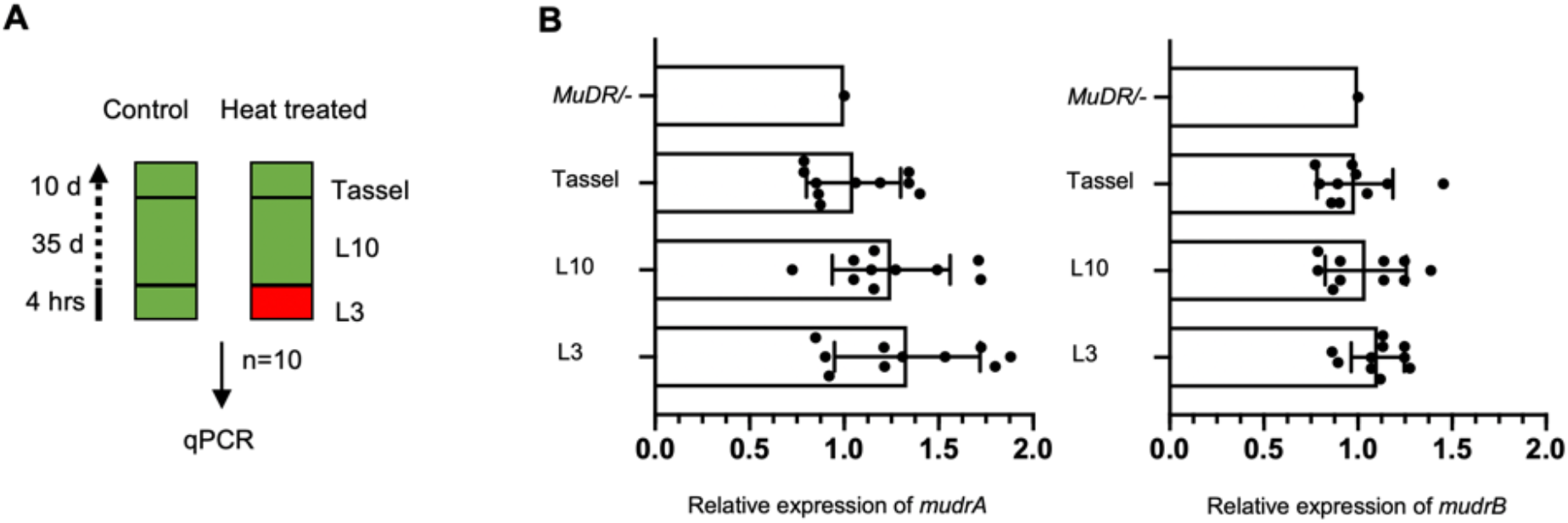
Expression of *mudrA* and *mudrB* in new emerging tissues following heat stress. (A) Diagram of the experiment. (B) qPCR was performed to measure transcript levels of *mudrA* and *mudrB* using expression of maize *Tub2* as an internal control. Expression levels were normalized to that of an active *MuDR* element, which was set at one. All data are the average of two technical replicates from ten independent plants. An unpaired t-test was performed. Error bars indicate mean ± standard deviation (SD) of the ten biological replicates.

### *MuDR* activity is stably heritably transmitted to subsequent generations

Our previous work had demonstrated that silenced *mudrA* (but not *mudrB*) can be progressively and heritably reactivated only after multiple generations of exposure to the *mop1* mutation under normal conditions. Only after eight generations could this activity could be stably transmitted to subsequent generations in the absence of the *mop1* mutation [35]. To determine if the somatic activity we observed after heat stress can be transmitted to the next generation, we crossed the heat-treated *mop1* homozygous plants that carried transcriptionally reactivated *MuDR* (designated *MuDR*^~^) and the sibling *mop1* homozygous *MuDR\** control plants, to a tester that was homozygous wild type for *mop1* and that lacked *MuDR* (Fig S1). MURA, the protein encoded by *mudrA* causes excision of a reporter element at the *a1-mum2* allele of the *A1* gene, resulting pale kernels with spots of colored revertant tissue. All plants used in these experiments were homozygous for *a1-mum2*. If *mudrA* were fully heritably reactivated, a cross between a *MuDR~/−; mop1/mop1* plant and a tester would be expected to give rise to 50% spotted kernels, and this phenotype would be expected to cosegregate with the reactivated *MuDR* element. The progeny of ten independent heat-reactivated individuals gave a total of 45% spotted kernels. In contrast, ten *mop1* homozygous siblings that carried *MuDR\** and that had not been heat treated gave rise to an average of only 0.7% spotted kernels after test crossing (Fig 6B, Supplemental Table 1). These results show that *MuDR* activity induced by heat treatment was transmitted to the next generation. RT-PCR in both endosperms and embryos of the spotted and pale progeny kernels and genotyping for the presence or absence of *MuDR* at position 1 on chromosome 9L [74] demonstrated that activity was transmitted to both the embryo and the endosperm, and that this activity cosegregated with the single *MuDR* present in these families (Fig S3). We employed a similar strategy to test stability of heritability. We crossed three subsequent generations to testers and counted the spotted kernels. We observed that the progeny of heat-reactivated individuals gave a total of 51%, 48% and 47% spotted kernels in the three subsequent generations. In contrast, subsequent generations of the lineage carrying *MuDR\** that had not been heat treated gave rise to only a small number of weakly spotted kernels (Fig 6C, D, Supplemental Table 1). These results demonstrate that heat reactivation is stable over multiple generations in a non-mutant genetic background, as is silencing in the absence of heat stress.

**Figure 6.**
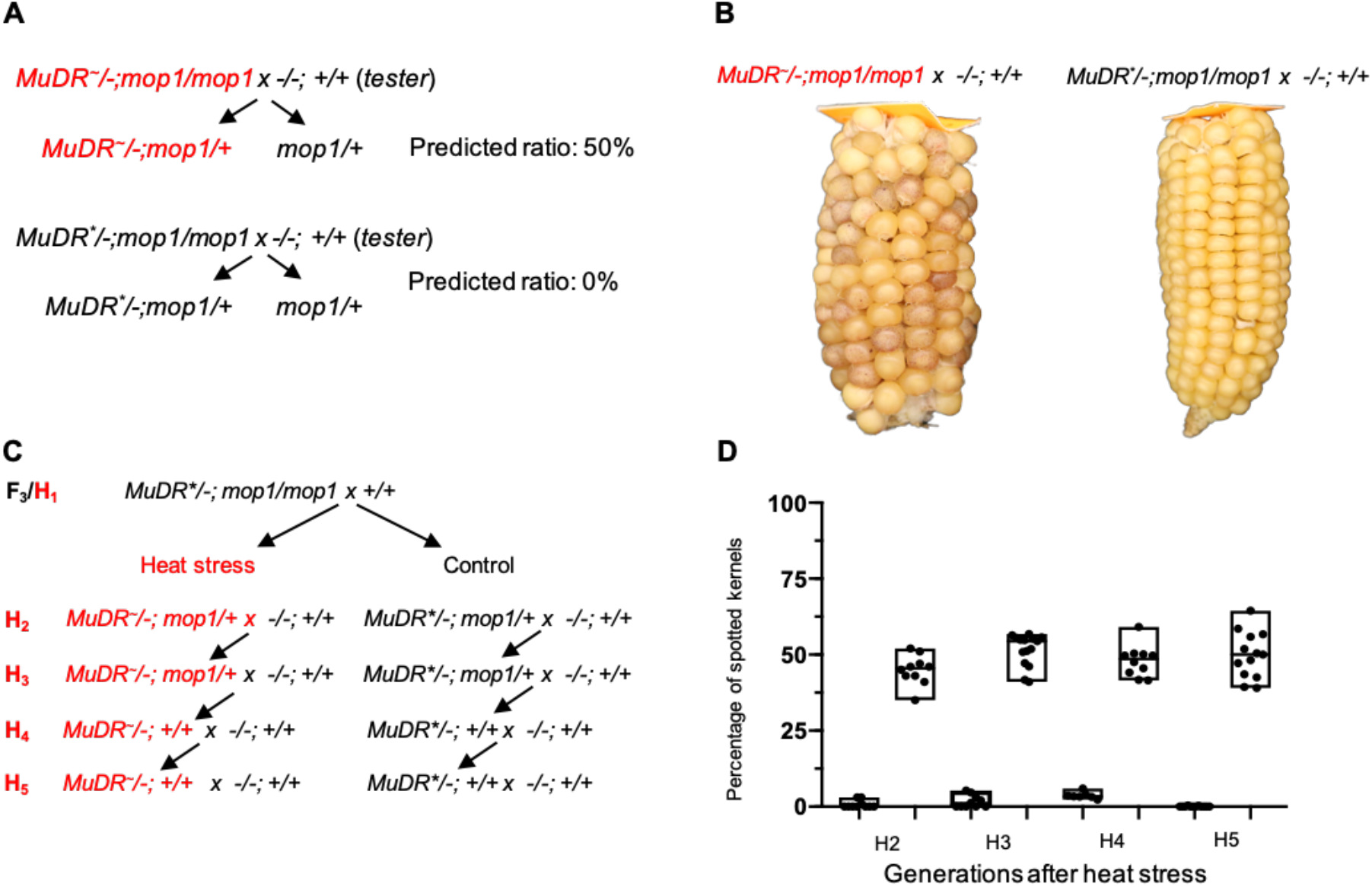
Testing transgenerational inheritance. (A) A schematic diagram showing the crosses used to determine transgenerational inheritance. (B) Ear ears derived from heat treated and control individuals. (C) Ratios of spotted kernels in generations of wild type plants following the heat stress (H1) generation.

### DNA hypomethylation is not associated with transgenerational inheritance of activity

We have shown that DNA methylation is not reduced under heat stress at TIRB, and that even a complete absence of methylation of TIRA under normal conditions does not result in transcriptional activation. These results suggest that, at least under normal conditions, DNA methylation of *MuDR* is neither necessary nor sufficient to mediate silencing. However, only plants that were *mop1* mutant and whose TIRs were missing either methylation of cytosines in all sequence contexts in the case of TIRA or those in the CHH sequence context in the case of TIRB were reactivated under heat stress. This suggests that a loss of methylation may be a precondition for initiation, and perhaps propagation, of continued activity after that stress. To test the later possibility, we examined DNA methylation at TIRA and TIRB in the *mop1* heterozygous H2 progenies of heat-reactivated *mop1* mutant plants and those of their unheated *mop1* mutant sibling controls (Fig S1). Surprisingly, we found that both TIRA and TIRB were extensively methylated in all three sequence contexts in all progeny examined regardless of their activity status (Fig 7). Indeed, their methylation was indistinguishable from that observed at silenced *MuDR* elements. This suggests that after reactivation, although the restoration of MOP1 does result in the restoration of methylation at both TIRA and TIRB, this methylation is not sufficient for reestablishment of silencing at either of these TIRs. In order to determine whether DNA methylation we observed in these wild type H2 plants was stable, we examined TIRA and TIRB methylation in plants four generation removed from the initial heat stress. Surprisingly, we found that the observed patterns of methylation in this generation at both TIRs closely resembled that of fully active *MuDR* elements (Fig 7). This suggests that patterns of methylation consistent with activity are in fact restored in the heat stressed lineage, but only after multiple rounds of meiosis in a non-mutant genetic background.

**Figure 7.**
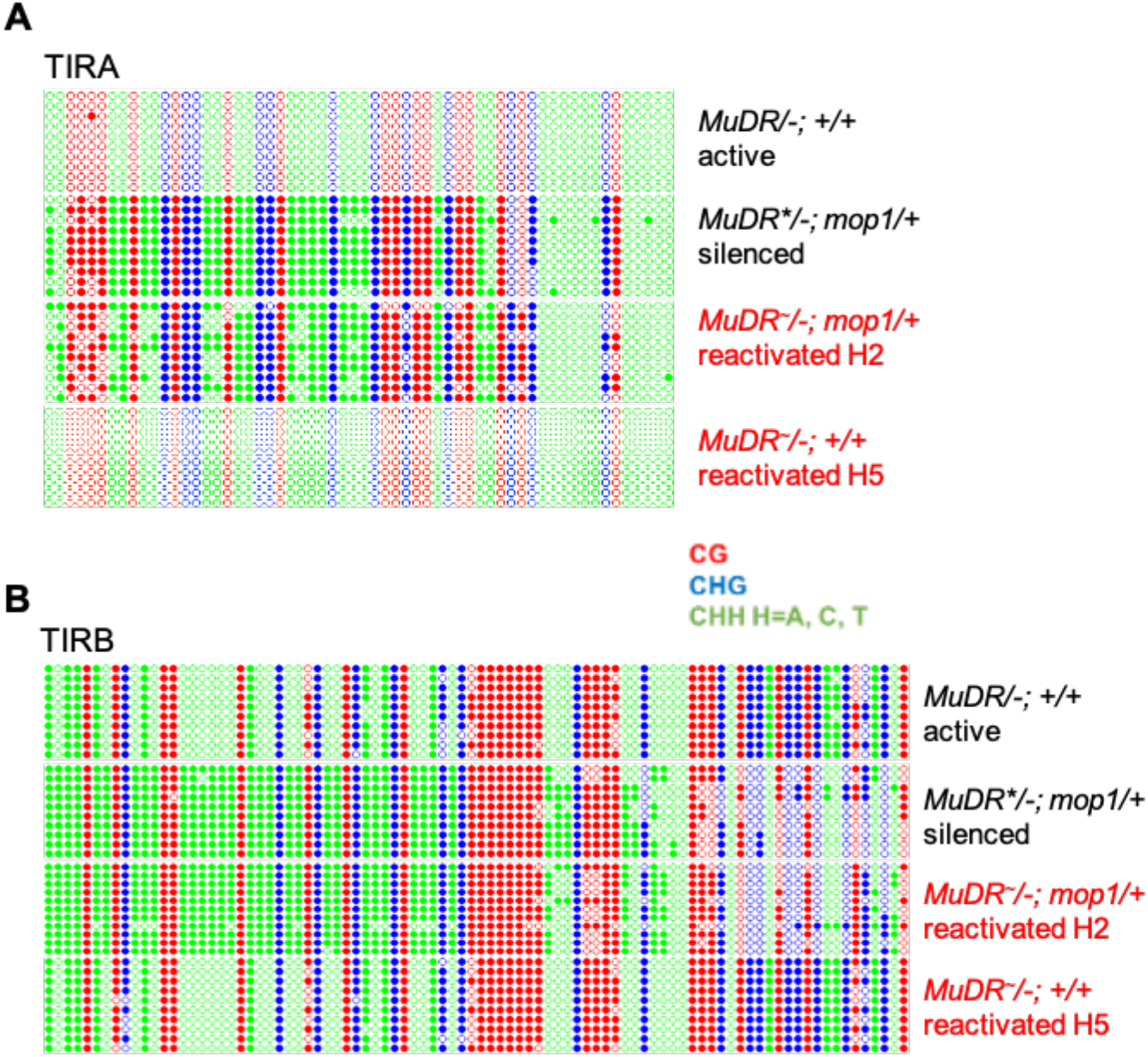
DNA methylation patterns at TIRA and TIRB of progeny of heat-treated H_2_ and H_5_ plants. (A) DNA methylation patterns at TIRA. (B) DNA methylation patterns at TIRB. Ten individual clones were sequenced from each amplification of bisulfite-treated sample. The cytosines in different sequence contexts are represented by different colors (red, CG; blue, CHG; green, CHH, where H=A, C, or T). For each assay, six independent samples were pooled together.

### Transgenerational heritability of activity is associated with heritability of histone modifications

DNA hypomethylation is not associated with transgenerational inheritance of *MuDR* activity, and DNA hypermethylation does not result in a restoration of silencing in wild type progeny of heat reactivated mutants. A plausible alternative is that the observed changes in histone marks mediate heritable propagation of activity of both *mudrA* and *mudrB* independent of methylation status. To test this hypothesis, we determined the levels of H3K9me2, H3K27me3 and H3K4me3 at TIRA and TIRB in the *mop1* heterozygous H2 progenies of heat-reactivated *MuDR~/−; mop1/mop1* plants and those of their sibling untreated *MuDR*/−; mop1/mop1* sibling controls. Consistent with the continued activity of *mudrB* in the progeny of the heat stressed plants, relative levels of H3K27me3 levels remained low and H3K4me3 remained high at TIRB in these plants, suggesting that heritable propagation of H3K27me3 is responsible for that continued activity (Fig 8). Similarly, at TIRA, H3K9me2 remained low and H3K4me3 remained high in these progenies. Interestingly, the increase in DNA methylation in these *MuDR* active *mop1* heterozygous plants was associated with a further decrease in levels of H3K9me2 at TIRA relative to that of their heat stressed *mop1* homozygous parents, down to the levels of the active *MuDR* control. This suggests that a increase in methylation of these active elements in the wild type background resulted in a concomitant decrease in H3K9me2 at TIRA.

**Figure 8.**
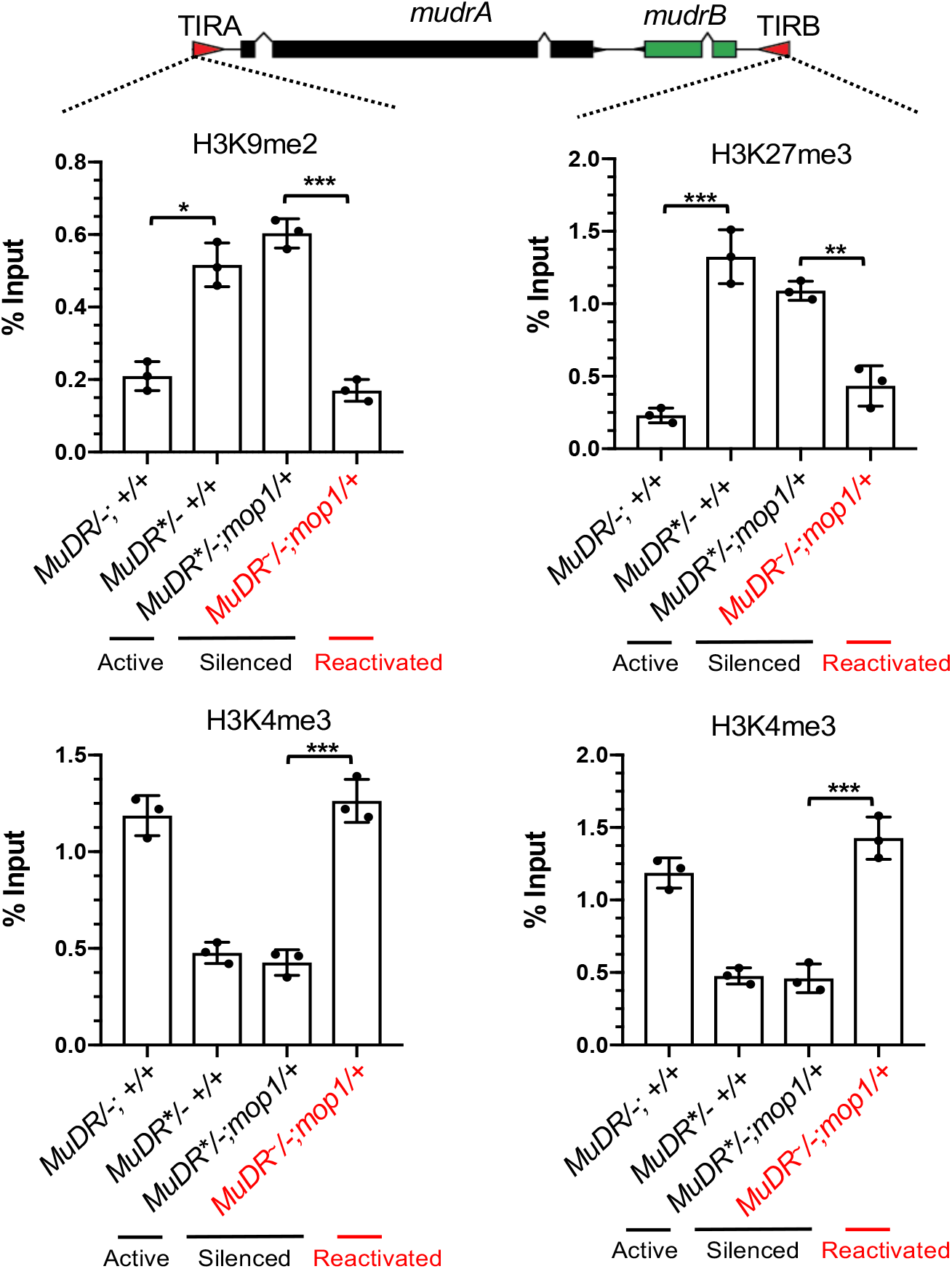
ChIP-qPCR analysis of enrichment of histone marks, H3K9me2, H3K27me3 and H3K4me3 at TIRA and TIRB. Relative enrichment of H3K9me2, H3K27me3 and H3K4me3 at TIRA and TIRB in leaf 3 of plants of the indicated genotypes. qPCR signals were normalized to *Copia* and then to the value of input samples. All data are the average of two technical replicates from three independent lines. An unpaired t-test was performed. Error bars indicate mean ± standard deviation (SD) of the three biological replicates. *P<0.05; **P < 0.01; ***P<0.001

## Discussion

### DNA methylation is neither necessary nor sufficient for the maintenance of silencing at TIRA or TIRB

Our results demonstrating that methylation is not necessary for maintenance of epigenetic silencing in *mop1* mutant plants (Fig 1) and is not sufficient to trigger silencing in H2 reactivated plants (Fig 7) suggest that at this particular locus, DNA methylation is not the key determinative factor with respect to either silencing or its reversal. In contrast, changes in H3K9me2 are closely correlated with changes in TIRA activity, suggesting that it is this modification, rather than DNA methylation, that mediates both activity and heritable transmission of silencing of *mudrA*. Given that H3K9me2 is normally tightly associated with cytosine methylation, particularly in the CHG context [21, 82], this result is unexpected. However, our results clearly demonstrate that this modification can be heritably propagated in the absence of DNA methylation and in the absence of the original trigger for silencing, *Muk*. Even more unexpected is our observation that, once *mudrA* becomes silenced, in *mop1* mutants there appears to be reciprocal relationship between DNA methylation of TIRA and H3K9me2 enrichment. Based on previous experiments, our expectation was that *mop1* would eliminate cytosine methylation in the 5’ end of TIRA, which is unrelated to transcriptional gene silencing of *mudrA*, but that it would not elimination of DNA methylation in the 3’ portion of TIRA, which is primarily in the CG and CHG contexts and is specifically associated with silencing of this gene [76]. In fact, we find that methylation in all three sequence context is eliminated throughout TIRA in *mop1* mutants, but this does not result in reactivation of *mudrA*. Instead, H3K9me2 actually significantly *increases* in the *mop1* mutant. This suggests that silencing at this locus is maintained via a balance between DNA and histone methylation, such that a loss of DNA methylation actually triggers an increase in histone modification. This in turn suggests that the state of activity of *mudrA* in some way determines the balance between histone and DNA modification, since neither modification by itself appears to be determinative. Our heat experiment supports this hypothesis. Heat rapidly reduces histone modification, but only back down to the level of the silent *mop1* heterozygous siblings rather that to the level of TIRA in an active element. In this case, the combination of an absence of DNA methylation with this reduced level of H3K9me2 appears to be sufficient to permit transcription of *mudrA*, as well as somatic propagation of the reactivated state to daughter cells after the heat is removed. Also supporting a balance hypothesis is the observation that in reactivated *mop1* heterozygous progeny of *mop1* homozygous heat treated plants, methylation is restored to that observed in silenced elements and levels of H3K9 dimethylation are then reduced to the level observed in active elements. This suggests that, again, levels of DNA and histone modification balance each other, such that in increase in methylation in the wild type progeny of reactivated *mop1* mutant plants results in a concomitant decrease in histone modification. Interestingly, however, after multiple generations in a wild type background, methylation levels are reduced to those of active *MuDR* elements, suggesting that this reduced methylation level is a consequence, rather than a cause, of maintenance of activity. Collectively, these data suggest that DNA methylation can be a lagging indicator that is responding to a given epigenetic state, rather than determining it.

There are other instances in which silencing can be reversed without a loss of methylation. For instance, mutations in the putative chromatin remodeler MOTHER OF MORPHEOUS1(MOM1) can result in activation of silenced transgenes and some endogenous loci in the absence of a loss of DNA methylation [83–85]. Similarly, Microrchidia (MORC) ATPase genes, as well the H3K27 monomethyltransferases ATXR5 and ATXR6 in Arabidopsis, are required for heterochromatin condensation and TE silencing but not for DNA methylation or histone modification associated with that silencing [86–88]. However, unlike reactivated *MuDR* elements in our experiments, reintroduction of the wild type MOM1 or MORC alleles result in immediate re-silencing. Finally, mutations in two closely related Arabidopsis genes, MAIL1 and MAIN, can also result in activation of a subset of Arabidopsis TEs in the absence of a loss of methylation [89].

### The RdDM pathway buffers the effects of heat stress on silenced *MuDR* elements

Heat stress rapidly reverses silencing and is associated with a reduction of H3K9me2, but only in a *mop1* mutant background. This suggests that although DNA methylation is not required for the maintenance of silencing of *mudrA* and is not sufficient to trigger *de novo* silencing of this gene, it is required to prevent a response to heat stress. Thus, we suggest that the primary role of DNA methylation in this instance is to buffer the effects of heat. We note that this observation is similar but distinct from what has been observed for the Onsen retrotransposon in Arabidopsis. In that case, although heat stress by itself can induce transcription of Onsen [9, 90], it is only when the RdDM pathway is deficient that new insertions are transmitted to the next generation. However in wild type progenies of heat stressed mutants, Onsen elements are rapidly re-silenced [91]. In contrast, reactivated *MuDR* elements remain active for at least five generations, despite the fact that the RdDM pathway rapidly restores DNA methylation at both TIRA and TIRB. This is likely due to differences between these two elements with respect to the means by which the two elements are maintained in a silenced state. In the absence of *Muk, MuDR* elements are stably active over multiple generations [74, 92]. This suggests that silencing of *MuDR* requires aberrant transcripts that are distinct from those produced by *MuDR* that are not present in the minimal Mutator line. Experiments involving some low copy number elements in Arabidopsis that are activated in the DNA methylation deficient *ddm1* mutant background suggest that the same is true for these elements as well; once activated, these elements remain active even in wild type progeny plants [93]. In contrast, evidence from other TEs suggests that transcripts from these elements or their derivatives contribute to their own silencing [39, 94, 95].

### Heritably transmitted silencing of TIRB is associated with H3K27me3

Our observation that transgenerationally heritable silencing of *mudrB* is associated with H3K27me3 was surprising, given that this mark is generally associated with somatic silencing of genes that is reset each generation [96]. However, in the absence of that resetting, silencing can be heritably transmitted to the next generation [50, 52]. Our data clearly shows that this is the case for *mudrB*, whose H3K27me3 enrichment can be heritably transmitted following the loss of *Mu killer* through at least two rounds of meiosis, and we have evidence that *mudrB* remains stably silenced for at least eight generations [35]. Given that there is no selective pressure to reset TE silencing mediated by H3K27me3, this is not surprising.

There is evidence that heat stress can heritably reverse H3K27me3 at specific loci. H3K27 trimethylation can be reversed by the H3K27me3 demethylase REF6, which acts in conjunction the chromatin remodeler BRAHMA (BRM) to relax silencing at loci containing CTCTGYTY motifs [97]. In Arabidopsis, under heat stress, HEAT SHOCK TRANSCRIPTION FACTOR A2 (HSFA2) activates REF6, which can in turn de-repress HSFA2 by reducing H3K27me3 at this gene. This feedback loop can extend to the progeny of heat stressed plants, resulting in a heritable reduction in levels of H3K27me3 at REF6 target genes [98, 99]. However, as in the case for all transgenerational shifts in gene expression, the effect is temporary, and both H3K27me3 and gene expression levels are restored to their original state after two generations.

### Conclusions

There is a growing body of evidence suggesting that whatever else they do, all silencing pathways can and do silence TEs, and in many cases may have evolved to do so. For instance H3K27me3 is largely associated with gene rather than TE silencing in higher plants, the bryophyte *Marchantia polymorpha*, which diverged from extant land plants 400 mya, appears to employ H3K27me3 as a mark for a substantial fraction of its heterochromatin, in place of H3K9me2 [100]. Similarly, a majority of silenced maternal copies of paternally expressed genes in Arabidopsis are marked by H3K27me3 in addition to H3K9me2 and DNA methylation [101]. There is also evidence that the original, ancestral role of H3K27me3 may be in TE regulation. In the single celled ciliate, *Paramecium tetraurelia*, loss of function of the Enhancer-of-zeste-like protein Ezl1, which can catalyze methylation of both H3K9 and H3K27, results in global de-repression of TEs with minimal effects on gene expression [102]. In multicellular organisms, epigenetic silencing of cell lineages via this pathway simplifies the problem of differentiation by heritably silencing whole suites of genes in tissues in which they are not needed. Single celled organisms do not have that requirement, but, like all other organisms, they do have a requirement to heritably silence TEs.

Overall, our data suggests that even when examining a single TE in a single organism, a wide variety of epigenetic processes can be seen to play a role in both silencing and its reversal. At TIRA, a loss of DNA methylation in *mop1* mutants is associated with what appears to be a compensatory increase in H3K9me2, which is heritably reversed by a brief exposure to heat. Heritable transmission of a reactivated state of *mudrA* is refractive to a restoration of DNA methylation, which instead appears to adjust over time to reflect that activity rather than to block it. In contrast to *mudrA* (and most other TE genes) heritable *mudrB* silencing is associated with H3K37me3 enrichment, which, like H3K9me2 enrichment at TIRA, is readily and heritably reversed by heat treatment. At both TIRA and TIRB, methylation is neither necessary nor sufficient for silencing, but a lack of MOP1 and an associated loss of DNA methylation at both TIRs does appear to be required to precondition both *mudrA* and *mudrB* for responsiveness to heat, consistent with a role for RdDM in buffering the effects of high temperature in maize. Clearly, these results are primarily phenomenological, as the precise mechanism for the reversal of silencing we observe remains a mystery. However, they do suggest that there is a great deal that we do not yet understanding about how silenced states can be maintained and how they can be reversed.

## Materials and Methods

### Plant materials

Maize seedlings and adult plants were grown in MetroMix under standard long-day greenhouse conditions at 26°C unless otherwise noted. The minimal Mutator line consists of one full-length functional *MuDR* element and one nonautonomous *Mutator* element, *Mu1. Mu killer* (*Muk*), a derivative version of the *MuDR* transposon, can heritably trigger epigenetic silencing of that transposon. *Mutator* activity is monitored in seeds via excisions of a *Mu1* element inserted into the *a1-mum2* allele of the *A1* gene, resulting in small sectors of revertant tissue, or spots, in the kernels when activity is present. When *MuDR* activity is absent, the kernels are pale. All plants described in these experiments are homozygous for *a1-mum2*. Although *MuDR* can be present in multiple copies, all of the experiments described here have a single copy of *MuDR* at position 1 on chromosome 2L [92].

All of the crosses used to generate the materials examined in this paper are depicted in Fig S1. Active *MuDR/−;mop1/mop1* plants were crossed to *Muk*/−;*mop1*/+ plants. The resulting progeny plants were genotyped to screen for plants that carried *MuDR, Muk* and that were homozygous for *mop1*, which were designated F_1_ plants. F_1_ plants were then crossed to *mop1* heterozygotes. Progeny plants lacking *Muk* but carrying silenced *MuDR* elements, designated *MuDR\**, were designated F_2_ *MuDR\** progeny. F_2_ *MuDR\** progeny that were homozygous for *mop1* were crossed to *mop1* heterozygotes. The resulting F_3_ plants were genotyped for the presence of *MuDR*. These plants were either homozygous or heterozygous for in *mop1*. These F_3_ plants were those that were used for the heat stress experiments. H1 refers to the first generation of these F_3_ plants that were subjected to heat stress, with successive generations designated H_2_, H_3_, etc… *MuDR* was genotyped using primers Ex1 and RLTIR2. Because Ex1 is complementary to sequences flanking *MuDR* in these families, this primer combination is specific to the single *MuDR* element segregating in these families. *Muk* was genotyped using primers TIRAout and 12-4R3. The *mop1* mutation was genotyped using primers ZmRDR2F, ZmRDR2R and TIR6. All primer sequences are provided in Table S2.

### Tissue Sampling

Plants used in all experiments were genotyped individually. The visible portion of each developing leaf blade, when it was ≈10 cm, was harvested when it emerged from the leaf whorl. Only leaf blades of mature leaves were harvested. For the heat reactivation experiment, seedlings were grown at 26 °C for 14 days with a 12-12 light dark cycle. Seedlings were incubated at 42 °C for 4 hours and leaf 3 was harvested immediately after stress treatment. As a control, leaf 3 was also collected from sibling seedlings grown at 26 °C. For each genotype and treatment, 12 biological replicates were used, all of which were siblings. Samples were stored in −80 °C. After sample collection, all seedlings were transferred to a greenhouse at 26 °C. In order to determine if reactivation could be propagated to new emerging tissues, leaf 10 at a similar stage of development (~10 cm, as it emerged from the leaf whorl) and the immature tassel (~20 cm) were collected from each individual (Fig. 4A). To determine if the application of heat stress at a later stage of plant development can promote reactivation, an independent set of these seedlings from the same family were used. A similar strategy was employed. However, in this case, seedlings were heat stressed for 4 hours after the plants had grown 28 days at 26 °C. Leaf 7 was collected instead (Fig. 3B). For the bisulfite sequencing experiment, leaf 3 was collected from each individual, when it was ≈10 cm, as it emerged from the leaf whorl. In order to minimize potential variation among different individuals, leaves from 6 individuals with the same genotype and treatment were pooled together. For the ChIP assays, a total of ~ 2 g of leaves from leaf 3 of 6 sibling plants with the indicated genotypes was harvested. Three independent sets of these sample collections were colected and analyzed for each genotype and treatment. Leaf samples were fixed with 1% methanol-free formaldehyde and then stored in −80 °C.

### RNA isolation and RT-PCR analysis

Total RNA was extracted using TRIzol reagent (Invitrogen) and purified by Zymo Direct-zolTM RNA Miniprep Plus kit. 2 μl of total RNA was first loaded on a 1% agarose gel to check for good quality. Then, RNA was quantified by a NanoDropTM spectrophotometer (Thermo Fisher Scientific) and reverse transcribed using an oligo-dT primer and GoScriptTM Reverse Transcriptase (Promega). The resulting transcribed cDNA was amplified for 29 cycles with primers specific for the alanine aminotransferase (*Aat*) transcripts (Zm00001d014258) with an annealing temperature of 55 °C used as a control to ensure equal starting amounts of cDNA. Samples were then amplified for 32 cycles using the primers specific for *mudrA* and *mudrB* with an annealing temperature of 59°C for both primer pairs. PCR products were electrophoresed on a 1.2% agarose gel. Quantitative RT-PCR was performed by using SYBR Premix Ex TaqTM (TaKaRa Bio) on a ABI StepOnePlusTM Real-Time PCR thermocycler (Thermo Fisher Scientific) according to the manufacturer’s instructions. Expression of *ZmHsp90* (Zm00001d024903) shown in Fig S3 was measured using primers HSP90-qPCR_F and HSP90-qPCR_F. Relative expression values for all experiments were calculated based on the expression of the reference gene, *ZmTub2* (Zm00001d050716) using primers TUB2-qPCR_F and TUB2-qPCR_R and determined by using the comparative CT method. Sequences for all primers used for RT-PCR are available in Table S2.

### Genomic Bisulfite Sequencing

These experiments were performed as previously described [76]. In brief, genomic DNA was isolated and digested with RNase A (Thermo Fisher Scientific). 2 μl of this DNA was loaded on a 1% agarose gel to check for good quality and then quantified using a Qubit fluorometer (Thermo Fisher Scientific). 0.5-1 μg of genomic DNA from each genotype and treatment were used for bisulfite conversion. The EZ DNA Methylation-Gold kit (Zymo Research) was used to perform this conversion. Fragments from TIRA and TIRB were PCR-amplified using EpiMark Hot Start Taq DNA Polymerase (New England BioLabs). For TIRA, the first amplification was for 20 cycles using p1bis2f and TIRAbis2R with an annealing temperature of 48 °C, followed by re-amplification for 17 cycles using TIRAbis2R and TIRAmF6 with an annealing temperature of 50 °C. Amplicons from TIRB were amplified for 30 cycles using methy_TIRBF and methy_TIRBR with an annealing temperature of 55 °C. The resulting fragments were purified and cloned into pGEM^®^-T Easy Vector (Promega). Ligations and transformations were performed as directed by the manufacturer’s instructions. The resulting colonies were screened for the presence of insertions by performing a colony-based PCR using primers of pGEMF and pGEMTR with an annealing temperature of 52 °C. The sequences of all primers are provided in Table S1. Plasmid was extracted from positive colonies using the Zyppy Plasmid Kit (Zymo Research). Plasmid from at least 10 independent clones were sequenced at Purdue Genomics Core Facility. The sequences were analyzed using kismeth (http://katahdin.mssm.edu/kismeth/revpage.pl)[103].

### Chromatin Immunoprecipitation (ChIP)

The ChIP assay was performed as described previously with some modifications [104–106]. Briefly, leaf samples were treated with 1% methanol-free formaldehyde for 15 minutes under vacuum. Glycine was added to a final concentration of 125 mM, and incubation was continued for 5 additional minutes. Plant tissues were then washed with distilled water and homogenized in liquid nitrogen. Nuclei were isolated and resuspended in 1 mL nuclei lysis buffer (50 Mm tris-HCl pH8, 10 mM EDTA, 0.25% SDS, protease inhibitor). 50 μl of nuclei lysis was harvested for a quality check. DNA was sheared by sonication (BioruptorTM UCD-200 sonicator) sufficiently to produce 300 to 500 bp fragments. After centrifugation, the supernatants were diluted to a volume of 3 mL in dilution buffer (1.1% Triton X-100, 1.2mM EDTA, 16.7mM Tris-HCl pH8, 167mM NaCl). Each sample of supernatant was sufficient to make 6 immunoprecipitation (IP) reactions. Every 500 μl sample was precleared with 25 μl protein A/G magnetic beads (Thermo Fisher Scientific) for 1 hour at 4 °C. After the beads were removed using a magnet, the supernatant was removed to a new pre-chilled tube. 50 μl from each sample was used to check for sonication efficiency and set aside to serve as the 10% input control. Antibodies used were anti-H3K9me2 (Millipore), H3K27me2 (Millipore), H3K27me3 (Active Motif), H3K4me3 (Millipore) and H3KAc (Millipore). After incubation overnight with rotation at 4°C, 30 μl of protein A/G magnetic beads was added and incubation continued for 1.5 hours. The beads were then sequentially washed with 0.5 mL of the following: low salt wash buffer (20 mM Tris (pH 8), 150 mM NaCl, 0.1% (wt/vol) SDS, 1% (vol/vol) Triton X-100, 2 mM EDTA), high salt wash buffer (20 mM Tris (pH 8), 500 mM NaCl, 0.1% (wt/vol) SDS, 1% (vol/vol) Triton X-100, 2 mM EDTA), LiCl wash buffer (10 mM Tris (pH 8), 250 mM LiCl, 1% (wt/vol) sodium deoxycholate, 1% (vol/vol) NP-40 substitute, 1 mM EDTA), TE wash buffer (10 mM Tris (pH 8), 1 mM EDTA). After the final wash, the beads were collected using a magnet and resuspended with 200 μl X-ChIP elution buffer (100 mM NaHCO3, 1% (wt/vol) SDS). A total of 20 μl 5M NaCl was then added to each tube including those samples used for quality checks. Cross-links were reversed by incubation at 65 °C for 6 hours. Residual protein was digested by incubating with 20 μg protease K (Thermo Fisher Scientific) at 55 °C for 1 hour, followed by phenol/chloroform/isoamyl alcohol extraction and DNA precipitation. Final precipitated DNA was dissolved in 50 μl TE. Quantitative RT-PCR was performed by using SYBR Premix Ex TaqTM (TaKaRa Bio) on an ABI StepOnePlusTM Real-Time PCR thermocycler (Thermo Fisher Scientific) according to the manufacturer’s instructions. The primers used in this study are listed in Table S2. The primers used to detect H3K9 and H3K27 dimethylation of Copia retrotransposons and H3K4 trimethylation of actin that were used as internal controls in this study have been validated previously [106]. Primers used for TIRA (TIRAR and TIRAUTRR) and TIRB (Ex1 and RLTIR2) were those used previously to detect changes in chromatin at these TIRs [75]. Expression values were normalized to the input sample that had been collected earlier using the comparative CT method.

## Acknowledgements

We thank R. Keith Slotkin for critical reading of the manuscript and Anthony Canon for testing the stability of transgenerational heritability.

## Supplemental Figures

**Fig S1.**
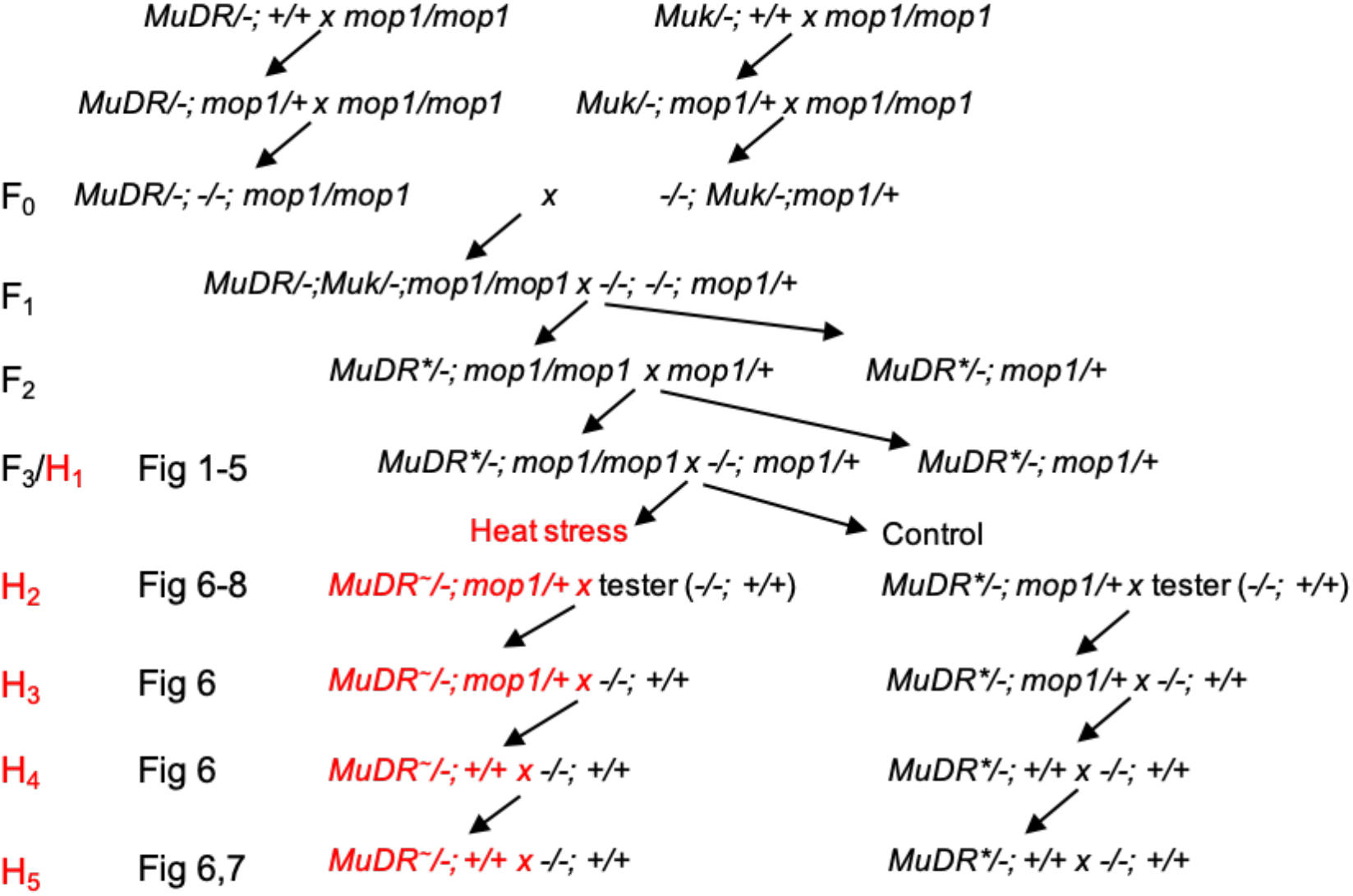
Diagram of the crosses and generations used in this study. F1 refers to the first generation during which *MuDR* was exposed to *Muk*. H_1_, which corresponds to F_3_, is the generation in which a brief heat treatment was applied. *MuDR* indicates an active *MuDR* element. *MuDR\** indicates an inactive *MuDR* element. *MuDR~* indicates a reactivated *MuDR* element.

**Fig S2.**
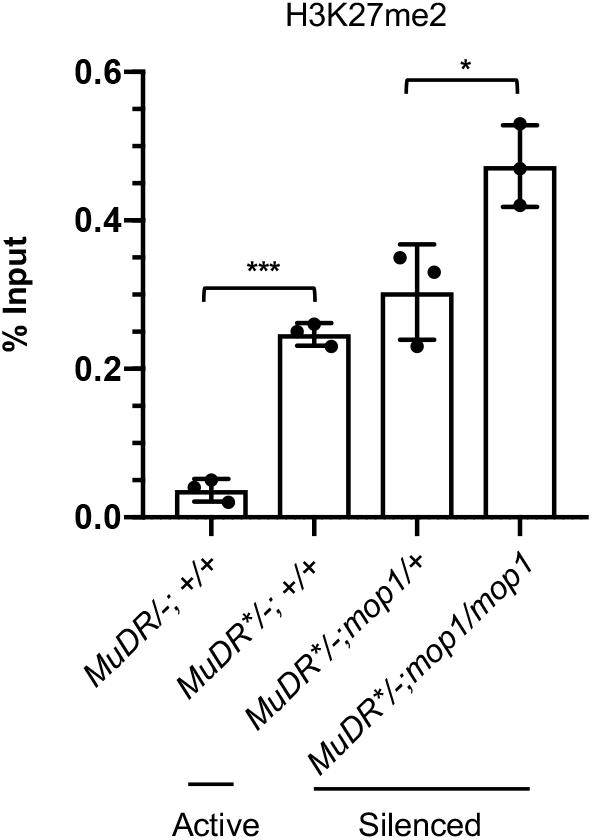
ChIP-qPCR analysis of enrichment of H3K27me2 at TIRA. Relative enrichment of H3K27me2 at TIRA in leaf 3 of plants of the4 indicated genotypes. The qPCR values were normalized to *Copia* and then to the value of input samples. All data are the average of two technical replicates from three independent sibling plants. An unpaired t-test was performed. Error bars indicate mean ± standard deviation (SD) of the three biological replicates. *P<0.05; ***P<0.001

**Fig S3.**
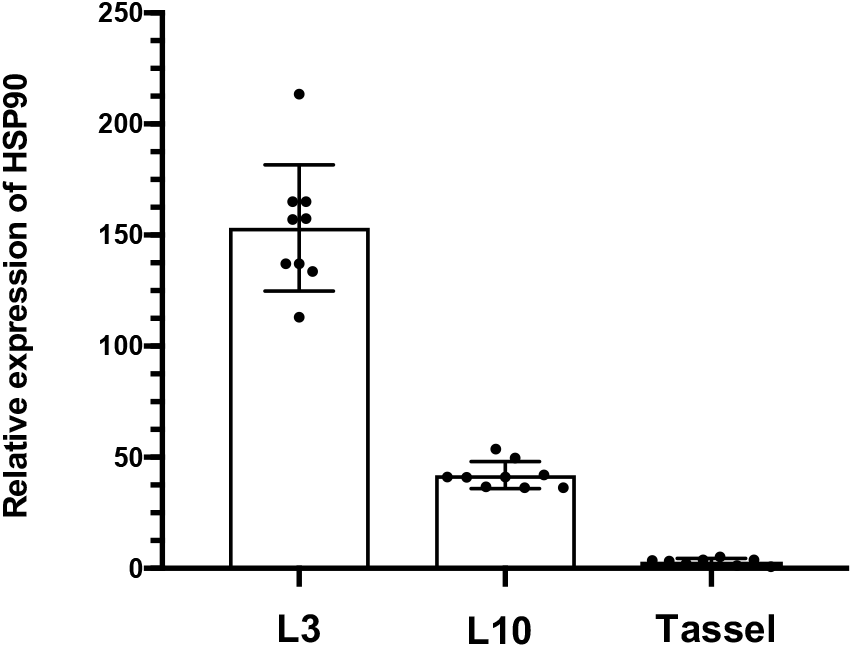
Real-time PCR analysis of *Hsp90* expression in the indicated tissues. Quantitative real-time PCR was performed to measure transcript levels of *ZmHsp90*. Data are the average of two technical replicates collected from ten independent lines. Error bars indicate mean ± standard deviation (SD) of the ten biological replicates.

**Fig S4.**
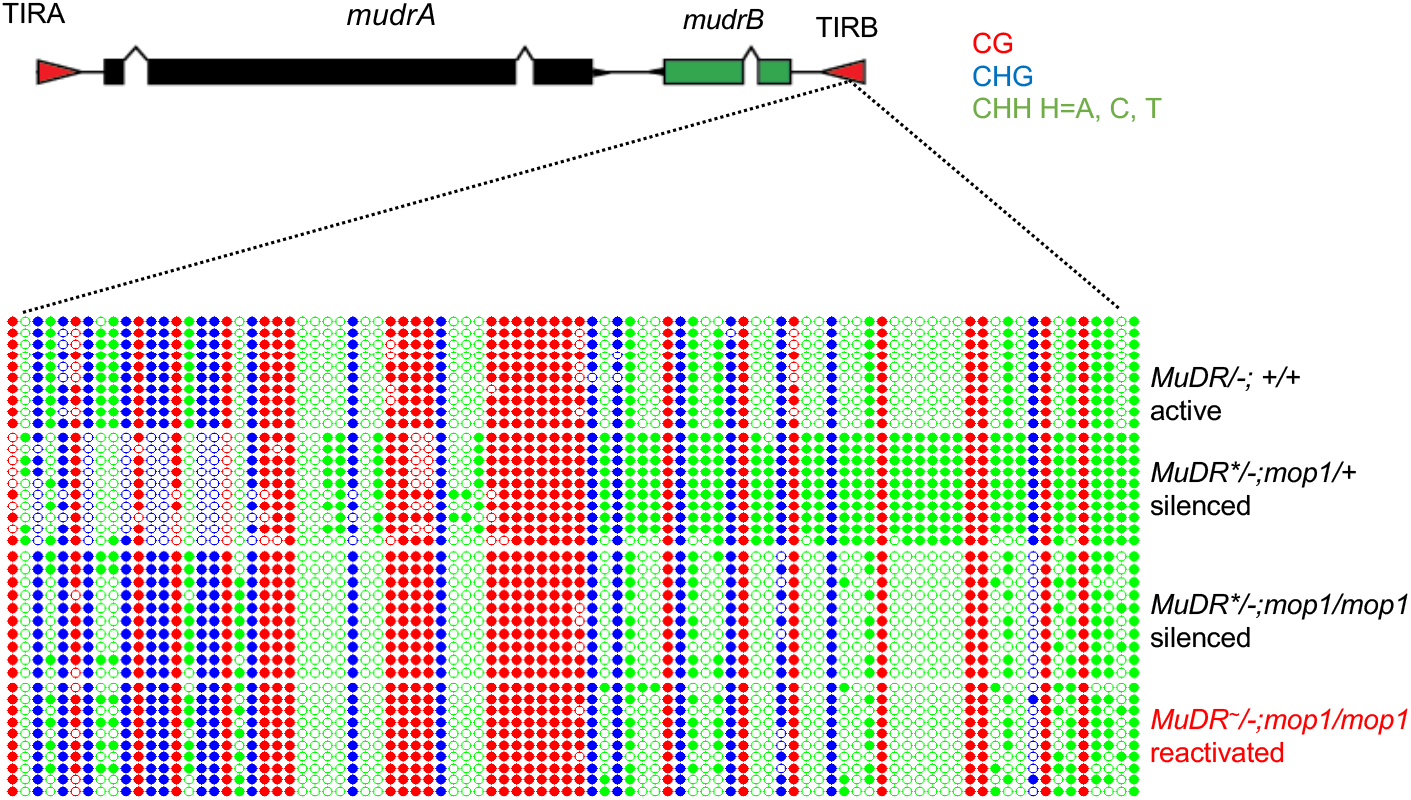
DNA methylation patterns at TIRB of heat-treated H1 *mop1mop1* plants. DNA methylation patterns at TIRA and TIRB. Ten individual clones were sequenced from each amplification of bisulfite-treated samples with the indicated genotypes. The cytosines in different sequence contexts are represented by different colors (red, CG; blue, CHG; green, CHH, where H=A, C, or T). For each sample, six independent samples were pooled together.

**Fig S5.**
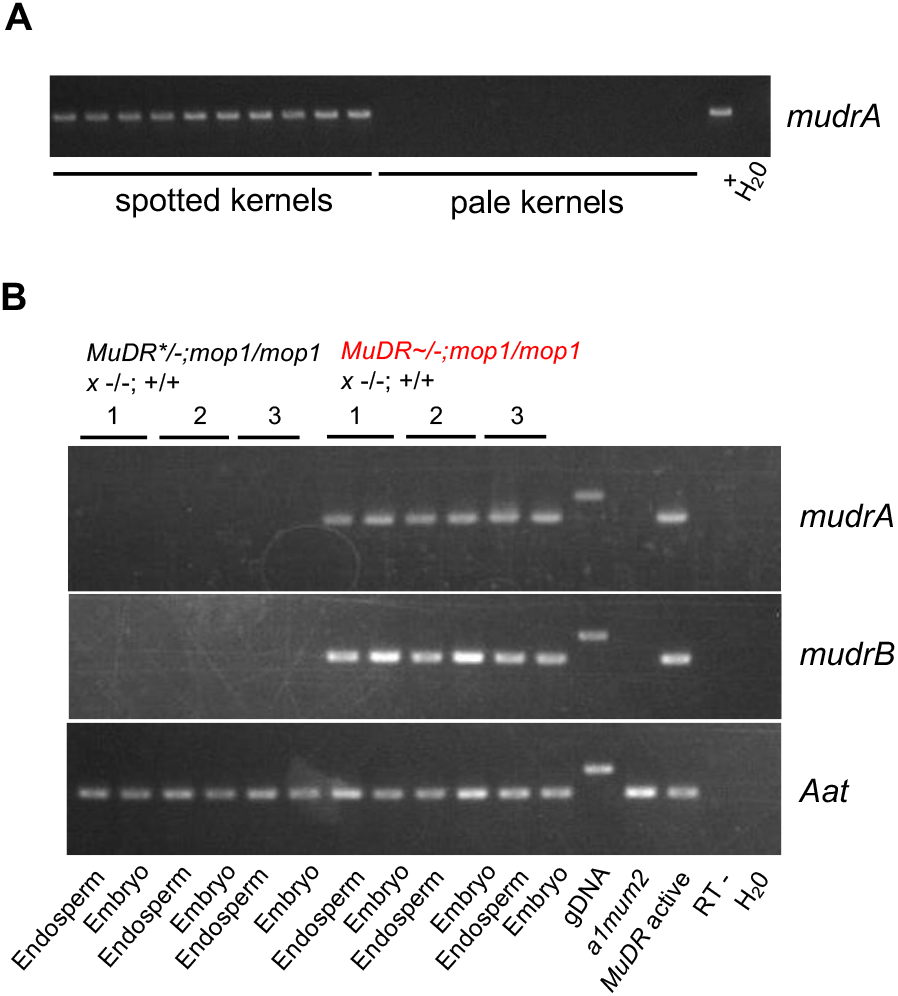
Analysis of *mudrA* and *mudrB* expression in progenies of H1 heat stressed plants. (A) Genotyping results of an ear from the H2 generation. (B) RT-PCR analysis of *mudrA* and *mudrB* expression in embryos and endosperms from kernels derived from three independent ears derived from crosses of H1 heat stressed plants and control siblings *Aat* is a housekeeping gene that serves as a positive control.

